# Late Bayesian inference in sensorimotor behavior

**DOI:** 10.1101/130062

**Authors:** Evan Remington, Mehrdad Jazayeri

## Abstract

Sensorimotor skills rely on performing noisy sensorimotor computations on noisy sensory measurements. Bayesian models suggest that humans compensate for measurement noise and reduce behavioral variability by biasing perception toward prior expectations. Whether the same holds for noise in sensorimotor computations is not known. Testing human subjects in tasks with different levels of sensorimotor complexity, we found a similar bias-variance tradeoff associated with increased sensorimotor noise. This result was accurately captured by a model which implements Bayesian inference after – not before – sensorimotor transformation. These results indicate that humans perform “late inference” downstream of sensorimotor computations rather than, or in addition to, “early inference” in the perceptual domain. The brain thus possesses internal models of noise in both sensory measurements and sensorimotor computations.

## Introduction

Consider the challenging task of returning a tennis serve. Not only must one accurately infer the path of the ball but also quickly transform that information into a motor plan that would yield a desirable outcome. The ability to apply such transformations is central to our behavioral repertoire and to the performance of athletes, musicians, and professionals such as surgeons and airplane pilots. It has been demonstrated that sensorimotor transformations are noisy (Soechting and Flanders 1989b; Pine et al. 1996; Gordon, Ghilardi, and Ghez 1994; McIntyre et al. 2000; Sober and Sabes 2005; Churchland, Afshar, and Shenoy 2006; Schlicht and Schrater 2007). The ubiquitous nature of sensorimotor transformations in behavior raises an important and unresolved question: does the brain have an internal model of sensorimotor noise (SMN), and do humans adopt strategies to mitigate its effects?

Research in the past several decades has tackled a similar question in the domain of sensory and motor systems, asking whether the brain is optimized to handle sensory and motor noise. Bayesian models have shown that humans adopt a number of strategies to minimize the effect of sensory and motor noise on behavior. For instance, when multiple sensory cues are available, humans rely more heavily on cues that are more reliable (R. J. van Beers, Sittig, and Gon 1999; Ernst and Banks 2002; Alais and Burr 2004; Bresciani, Dammeier, and Ernst 2008). Similarly, humans use their prior knowledge of statistics of sensory inputs to improve sensory estimates (Weiss, Simoncelli, and Adelson 2002; Körding and Wolpert 2004; Tassinari, Hudson, and Landy 2006; Jazayeri and Shadlen 2010). Optimal strategies in the presence of motor noise have also been reported, for example, when some movements are made more costly than others (Trommershäuser et al. 2005; Landy, Trommershäuser, and Daw 2012). These and related observations in multiple modalities (Battaglia, Jacobs, and Aslin 2003; Körding, Ku, and Wolpert 2004; Schlicht and Schrater 2007; Burge, Ernst, and Banks 2008; Butler et al. 2010) have provided strong evidence that the brain has an internal model of noisy representations in the sensory and motor systems and implements strategies to reduce their degrading effect on behavior.

Motivated by the success of normative models showing that the brain seeks to optimize behavior in the presence of sensory and motor noise, we hypothesized that the brain might additionally be equipped with mechanisms to minimize the effects of additional noise introduced by the transformation of sensory inputs to motor outputs (i.e., SMN). This question is particularly important as it bears on where in the brain Bayesian inferences are made (**Figure 1**). Since existing evidence supports perceptual Bayesian inference in the sensory domain (Weiss, Simoncelli, and Adelson 2002; Kersten, Mamassian, and Yuille 2004; Girshick, Landy, and Simoncelli 2011), an observation that Bayesian estimation does not take SMN into account would support the notion that observers employ an “early inference” strategy in which the inference is made within the sensory system before a sensorimotor transformation is applied. If, on the other hand, the brain takes SMN into account, we would conclude that the inference is made downstream after the sensorimotor transformation stage, thus providing evidence for a “late inference” strategy in the association and/or premotor brain areas. Critically, the early and late inference strategies make distinct predictions about the effect of SMN on behavior. With an early inference strategy, any increases in SMN would lead to comparable increases in behavioral variability. In contrast, in the late inference model, the brain would counter this variance by using knowledge about the distribution of responses (i.e., prior distribution). This strategy would introduce additional biases toward the mean of the prior distribution and lead to an overall improvement in performance.

**Figure 1.**
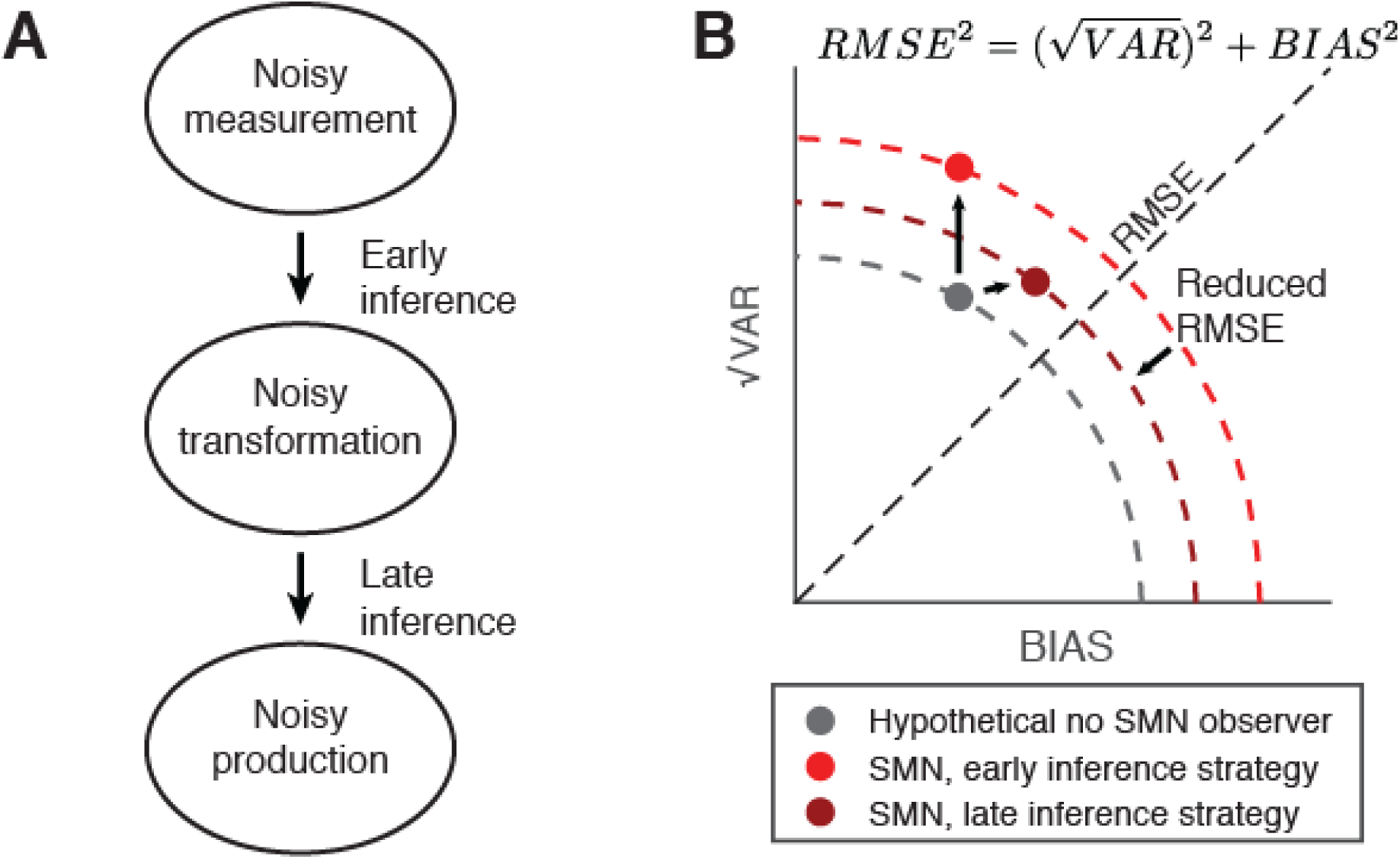
Bayesian inference in the presence of sensorimotor noise (SMN). **A**. A schematic of a sensorimotor task comprising sensory measurement, sensorimotor transformation, and motor production, all subject to internal noise. In sensorimotor tasks which don’t explicitly model SMN, subjects’ behavior is consistent with a strategy in which the effect of sensory noise (“noisy measurement”) on behavior is mitigated by Bayesian inference which biases perceptual estimates towards the mean of a sensory prior, reducing behavioral variability. We refer to this strategy as “early inference.” An alternate strategy, which we term “late inference” incorporates a prior on behavioral responses to mitigate the effects of both sensory and sensorimotor noise (“noisy transformation”). Motor noise (“noisy production”), such as that inherent in motor neurons and muscles, cannot be mitigated by relying on prior information. (**B**) An illustration of relationship between overall variability (*√VAR*), overall bias (*BIAS*), and *RMSE* for the early Inference and late Inference strategies in the presence of SMN. The equation shows the mathematical relationship between (*√VAR*), *BIAS*, and *RMSE*. This relationship can be depicted on a quarter circle (dashed lines) with the radius representing *RMSE*. The gray circle represents (*√VAR*) and *BIAS* values for hypothetical observer with sensory and motor noise only, and no sensorimotor noise (“no SMN observer”). For this case, early inference and late inference strategies produce identical behavior (gray). Introducing SMN will cause *RMSE* to increase. For an observer that uses an early inference strategy, the increase in *RMSE* will manifest primarily as increased *√VAR* (bright red). In contrast, the effect of SMN in an observer using a late inference strategy (dark red) would be primarily an increase in *BIAS*. Crucially, the late inference strategy would lead to a smaller increase in *RMSE*, as represented by the smaller radius of the dark circle compared to the bright one.

To distinguish between the early and late inference strategies, we exploited the observation that more complex sensorimotor transformations engender more noise (Soechting and Flanders 1989b; Pine et al. 1996; McIntyre et al. 2000; Sober and Sabes 2005; Schlicht and Schrater 2007). We devised two experiments: (1) a time interval estimation and production task, and (2) a length estimation and production task. In each task, we compared subject’s performance across two sensorimotor contexts. In one context, which we refer to as the “identity context”, the produced quantity (time interval or length) had to match a previously measured quantity. This was compared to a more complex “remapped context” in which subjects had to produce a quantity by applying a non-identity transformation to the sensory quantity. For example, subjects had to produce a length that was 50% longer than the stimulus.

As expected, the remapped context negatively impacted performance in both tasks, revealing the degrading effect of SMN. Importantly, increases in SMN in the remapped context led to increased biases toward the mean of the sensorimotor prior, an indication of the late Bayesian inference that takes SMN into account. These results reveal that the brain has an internal model of the noise associated with sensorimotor transformations and integrates this information with prior knowledge to optimize inferences in terms of produced quantities.

## Results

We conducted two psychophysical experiments, one involving measurement and production time intervals, and another involving measurement and production of lengths. Each trial in each task consisted of two epochs: a measurement epoch during which a sensory quantity (time interval or length) was measured, and a subsequent production epoch during which subjects had to produce a quantity based on the preceding measurement. For each task, performance was quantified in two sensorimotor contexts: an identity context in which the produced quantity had to match the sensory quantity, and a remapped context in which the produced quantity had to match the sensory quantity multiplied by a fixed scale factor.

## Time measurement and production task: Ready, Set, Go

Eight human subjects performed a time interval measurement and production task (**figure 2A**), also known as the “Ready, Set, Go” task, similar to a previous study (Jazayeri and Shadlen 2010). During the measurement epoch, subjects were presented with a sample interval (*t*_*s*_; see **Table 1** for all variables and abbreviations) demarcated by two visual flashes, “Ready” and “Set” (**figure 2A**). Subjects had to measure *t*_*s*_ and produce an interval (*t*_*p*_) afterwards by a key press (“Go” - no flash). The interval *t*_*p*_ was measured from the start of the Set flash until the key press. In both identity and remapped contexts, *t*_*s*_ was drawn from the same discrete uniform prior distribution with 11 values ranging from 600 and 1000 ms. In the identity context, the correct interval (*t*_*c*_) was the same as *t*_*s*_, and in the remapped context, *t*_*c*_ was 1.5 times *t*_*s*_. In other words, the two contexts were identical during the measurement epochs but differed with respect to the production epoch. We denote these two contexts in terms of a gain factors relating *t*_*c*_ to *t*_*s*_: gain = 1 for the identity context, and gain = 1.5 for the remapped context. Subjects received trial-by-trial feedback about their performance (see Methods).

**Table 1.** Variables and abbreviations.

|  |  |
| --- | --- |
| **BIAS** | Summary bias |
| $I_c$ | correct length interval |
| $I_s$ | sample length interval |
| **RMSE** | Root mean square error |
| $\sigma_p$ | Production standard deviation |
| **SMN** | Sensorimotor noise |
| $t_c$ | Correct time interval |
| $t_i$ | Inferred time interval |
| $t_m$ | Measured time interval |
| $t_p$ | Produced time interval |
| $t_s$ | Sample time interval |
| $\sqrt{\text{VAR}}$ | Summary standard deviation |
| $w_m$ | Measurement Weber fraction |
| $w_p$ | Production Weber fraction |
| $w_t$ | Sensorimotor transformation Weber fraction |
| **deg** | Visual degrees |
| **ms** | Milliseconds |
| $H_0$ | Null hypothesis |

We quantified performance with three statistics (Jazayeri and Shadlen 2010): *BIAS*, which summarizes the deviation of average responses from the correct interval, *√VAR*, which summarizes the variability of responses across *t*_*s*_, and *RMSE*, which summarizes the total root mean square error. The three quantities are related through a sum of squares: *RMSE*^2^ *=* (*√VAR*)^2^ *+ BIAS*^2^ (**Figure 1B**). To ensure that the results were not influenced by overall tendencies to be late or early for all intervals, we calculated these statistics after removing an offset term that accounted for subjects’ overall bias (see Methods). For most subjects, the offset term was relatively small (**see Supplementary Table 1**).

**Figure 2B** illustrates the behavior of a typical subject in the identity (gray) and remapped (red) contexts of the timing task. Subjects’ behavior in the identity context exhibited prior-dependent bias, consistent with Bayesian integration as was shown previously (Jazayeri and Shadlen 2010; Acerbi, Wolpert, and Vijayakumar 2012; Cicchini et al. 2012) (**figure 2B**, in gray). In the remapped context, subjects had to perform the same task but with a gain of 1.5. We hypothesized that this more challenging sensorimotor transformation would cause an increase in SMN, and would thus increase the total *RMSE*. Additionally, we hypothesized that the increase in *RMSE* would be predominantly due to an increase in bias, consistent with the late inference hypothesis.

**Figure 2.**
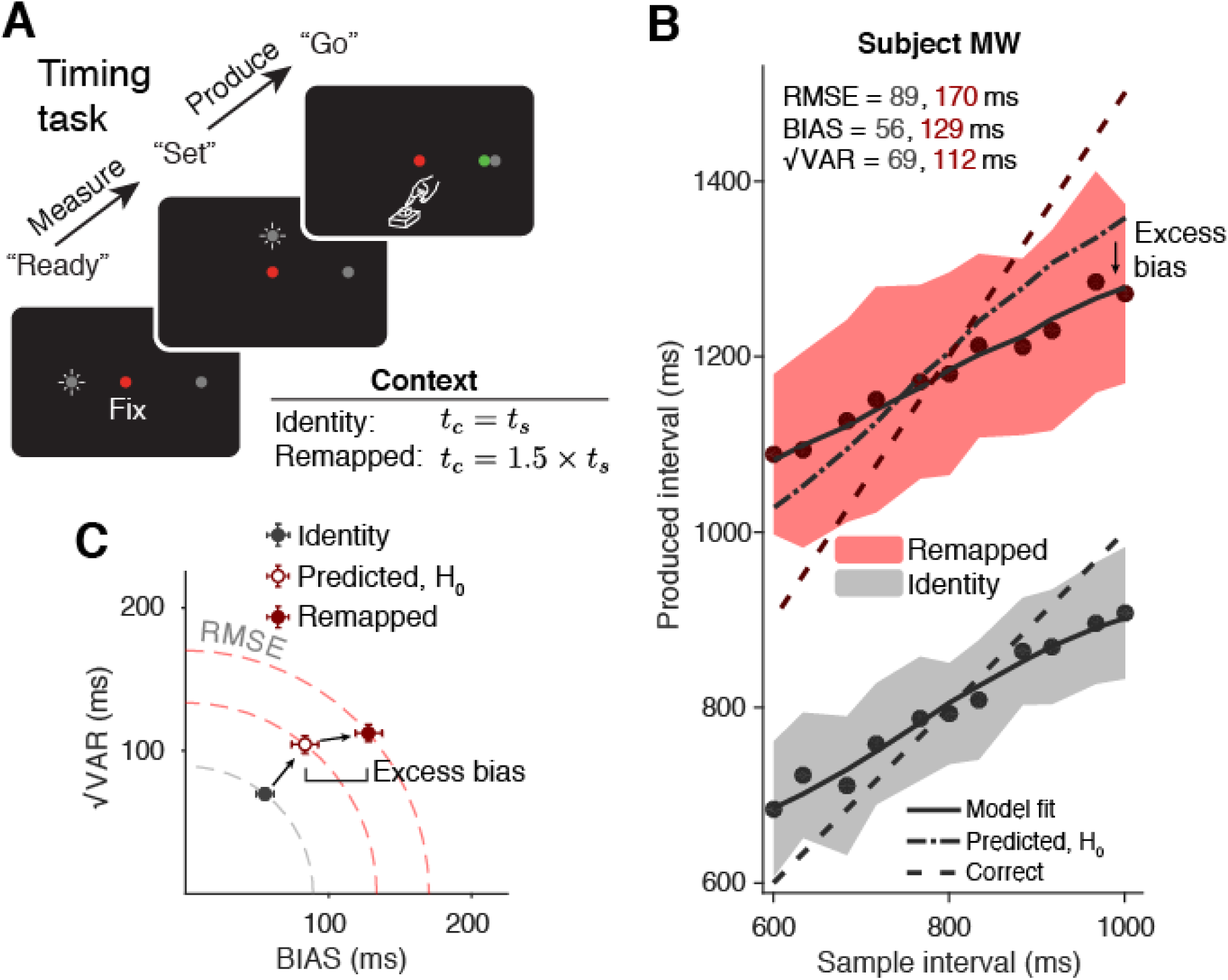
Time measurement and production task. **A**. Trial structure. Each trial began with the presentation of a red fixation spot. Subjects had to measure a sample time interval *t*_*s*_ demarcated by two flashes (“Ready” and “Set”) to the left of and above the fixation point, respectively. After Set, subjects pressed a key (“Go”) to produce an interval as close as possible to the correct interval *t*_*c*_ = *gain × t*_*s*_, where *gain* changed across two contexts. In the “identity” context, the correct interval was the same as *t*_*s*_ (*gain* = 1), whereas in the “remapped” context, the gain was 1.5. The position of a stimulus to the right of the fixation point served as a gain instruction cue. The distance of this stimulus to the fixation point is equal to or 1.5 times the distance of Ready to the fixation point for the gain of 1 and 1.5, respectively. After the response, subjects received scaled and signed feedback via the position of a colored circle (see Methods). **B**. Performance of an example subject in the identity (gray) and remapped (red) contexts. Filled circles and shaded regions indicate mean response times ± one standard deviation; dashed lines represent correct intervals. Solid lines represent the mean responses of a Bayesian observer model (see Methods) fit to the subject’s data separately for the two contexts; the dash-dot line in the *gain* = 1.5 condition corresponds to the prediction for the remapped context under the null hypothesis, using parameters of the model fit to the identity context (H_0_: no additional SMN). The subject’s behavior shows excess bias beyond what was predicted assuming no additional SMN. **C.** *√VAR* vs. *BIAS* for the two contexts (gray for identity, and dark red for remapped), as well as the prediction for the remapped context assuming no additional SMN (empty circle). This prediction underestimates *RMSE* indicating larger SMN for the remapped context. The increased RMSE was predominantly due to an increase in BIAS (“excess bias”). Dashed quarter circles illustrate combinations of *BIAS* vs. *√VAR* giving rise to equal *RMSE*; error bars are computed from the standard deviation of bootstrapped estimates (n = 1000).

We tested the first hypothesis (increased SMN in the remapped context) by comparing behavior in the remapped context to that predicted under the assumption of no additional SMN. This null hypothesis can be formulated straightforwardly by applying the gain factor of 1.5 to the estimates of *t*_*s*_ and taking into account the additional variability in *t*_*p*_ due to the linear scaling of production noise (Rakitin et al. 1998; Gallistel and Gibbon 2000). This leads to a simple prediction: without additional SMN, *RMSE*, *BIAS*, and *√VAR* in the remapped context should be 1.5 times their values in the identity context (**figure 2B,C**). As shown by the example subject (**figure 2C**) as well as results across all eight subjects (**figure 3A, top**), the observed *RMSE* in the remapped context was consistently and significantly higher than expected under the assumption of no additional SMN (RMSE median = 130 ms, interquartile range = 20 ms vs. median = 180 ms, IQR = 40 ms, p = 0.016, Wilcoxon signed-rank test). This provides direct evidence that SMN increased in the remapped context and validates the success of our experimental design in manipulating SMN independently of sensory noise.

Having established an increase in SMN in the remapped context, we tackled the second hypothesis of whether the *RMSE* increase was due to an increased bias, as would be predicted if subjects optimized their behavior to mitigate the effect of SMN with the late inference strategy. The null hypothesis, which states that subjects do not optimize their behavior in the presence of SMN, can be formulated by an early inference strategy. In this strategy, subjects take the sensory noise into account but ignore SMN. This early inference strategy predicts that the increase in *RMSE* in the remapped context should be explained by an increase in *√VAR* and not *BIAS*. This is because, in the early inference strategy, SMN is introduced after the inference stage and thus can only lead to increased variance (see **Figure 1B**). As shown in **Figure 2C** for one example subject, the increase in *RMSE* was largely due to an increase in *BIAS*, which can be readily seen as an excess bias compared to the no additional SMN prediction (**Figure 2B**, Excess bias). The results for all subjects, summarized in **Figure 3A,** indicate a clear increase in the *BIAS* for the remapped context relative to the null prediction (*BIAS* median = 73 ms, interquartile range (IQR) = 30 ms vs. median = 130 ms, p = 0.023). Across subjects, there was also a small but consistent effect on *√VAR* (median = 106 ms, IQR= 15 ms vs. median = 113 ms, IQR= 28 ms, p = 0.008). *RMSE*, *BIAS*, and *√VAR* for individual subjects are summarized in **Supplementary Table 1**. The substantial increase in *BIAS* across subjects rejects the null hypothesis and provides evidence for the late Bayesian inference, and indicates that humans take SMN into account to optimize their responses.

**Figure 3.**
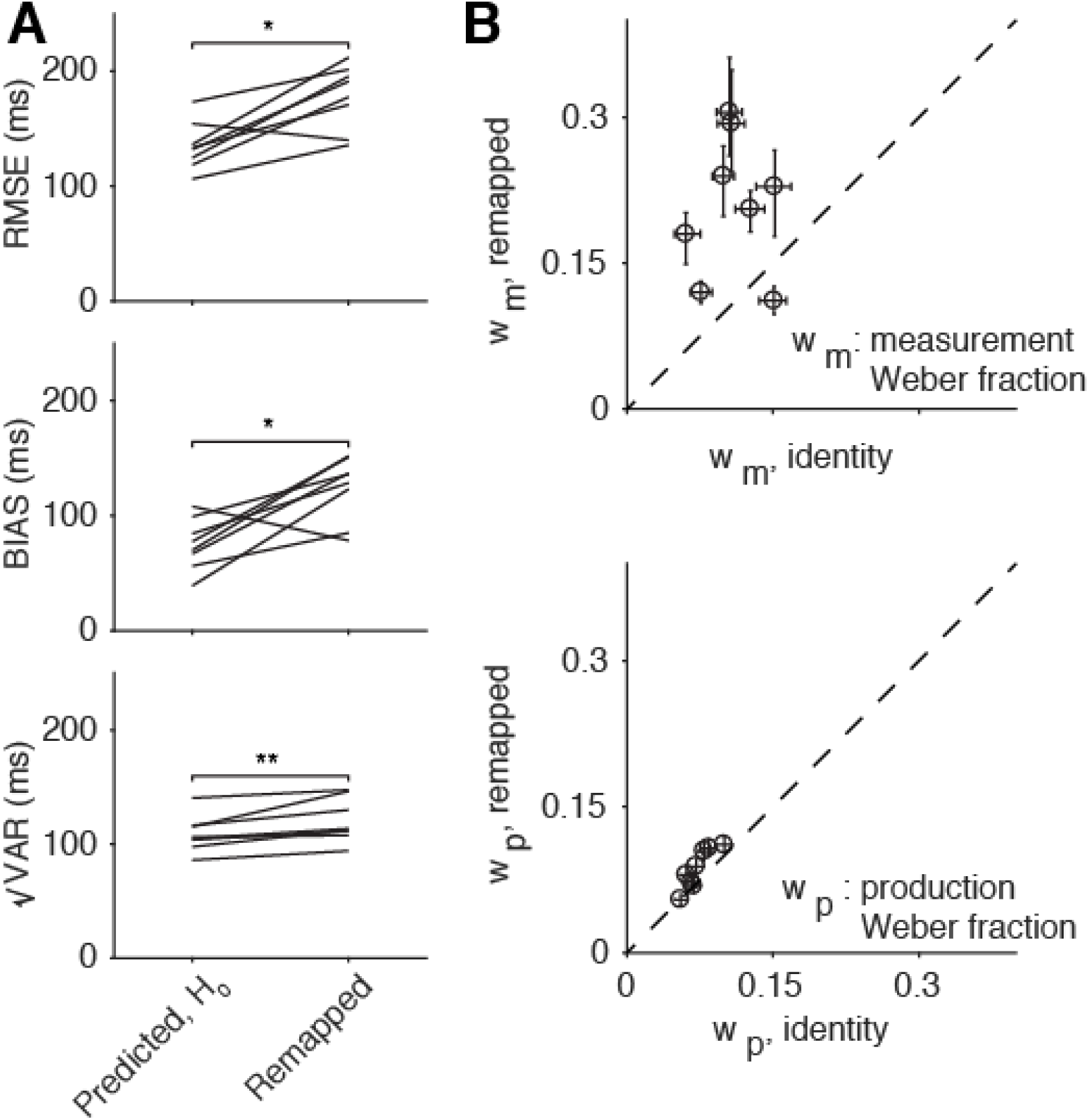
Summary of subjects’ behavior in the time measurement and production task. (**A**). Comparison of *RMSE* (top), *BIAS* (middle), and *√VAR* (bottom) for all subjects. The lines connect values predicted in the remapped context assuming no additional SMN (predicted, H_0_; left) based on the identity context (i.e. multiplied by the gain of 1.5) to actual values observed from behavior (*gain* = 1.5; right). Almost every subject had higher *RMSE* and *BIAS* than was predicted assuming no additional SMN. There was also a small increase in *√VAR* compared to predictions (*: p < 0.05, **: p < 0.01). (**B**) Parameters of observer model fits. We fit an observer model (see methods) to each subject’s data independently for the two contexts. This model did not explicitly account for SMN, and so any differences in SMN across contexts were reflected in the measurement and production Weber parameters (*w*_*m*_ and *w*_*p*_). Most subjects were fit with much higher values of *w*_*m*_ (top) in the remapped context, reflecting additional reliance on prior information consistent with a late inference strategy. There was also a more modest increase in *w*_*p*_ for the remapped context (bottom). Error bars represent 95% confidence intervals estimated using a bootstrap procedure (n = 1000).

Next, we compared the behavior of subjects in the two contexts using a Bayesian observer model (**Supplementary Figure 1**; Equation 4). This model comprises a noisy measurement stage (with scalar noise parameterized by *w*_*m*_), followed by a Bayesian estimation stage (Bayes-Least Squares), followed by a noisy production stage (with scalar noise parameter *w*_*p*_) (Jazayeri and Shadlen 2010). The model established Bayes-optimal behavior in the identity context (**Figure 2B**, “model fit,” bottom). We then fit the same model to subjects’ data in the remapped context allowing the parameters (*w*_*m*_ and *w*_*p*_) to take different values (**Figure 2B**, “model fit,” bottom). Without a mechanism to take SMN into account explicitly, the fits of this model to the remapped context would misattribute the drop in performance as being due to higher noise levels in the measurement (*w*_*m*_) and/or production (*w*_*p*_) relative to the identity context. As such, increases in *BIAS* would result in larger *w*_*m*_, and increases in *√VAR*, in larger *w*_*p*_ (**Supplementary Figure 2**). Since the early and late inference strategies are associated with increases in BIAS and *√VAR* respectively, we predicted that fitting this model to the data in the remapped context would result in a systematic increase in *w*_*m*_ and not *w*_*p*_. Model fits supported this prediction: *w*_*m*_ were substantially higher in the remapped context compared to the identity context (**Figure 3B**, also see **Supplementary Table 1**). This result further substantiates the hypothesis that increased SMN in the remapped context led to additional bias consistent with a late Bayesian inference strategy (**Figure 1A**).

To further validate this conclusion, we formulated a “Late-Inference” model in which we held *w*_*m*_ and *w*_*p*_ constant across contexts and added an additional scalar SMN parameterized by *w*_*t*_ in the remapped context. This model made late inference by virtue of the fact that inference was made after the introduction of *w*_*t*_. This Late-Inference model accurately captured the tradeoff between bias and variance in both contexts (**Figure 4A**), consistent with our hypothesis that additional bias in the remapped context was driven by increased SMN. We contrasted this model with two alternatives. First, we tested for the necessity of the additional scalar SMN by constructing an “Equal-SMN” model which omitted *w*_*t*_. This model was unable to simultaneously capture *RMSE*, *BIAS*, and *√VAR* in both contexts. Importantly, it systematically underestimated the bias in the remapped context (**Figure 4B**), validating the need for additional *w*_*t*_ in the remapped context. The second alternative model contrasted the Late-Inference model with an “Early-Inference” model in which the inference was made prior to the SMN. Similar to the Equal-SMN model, the Early-Inference model failed to capture behavior (**Figure 4B**), highlighting the importance of late inference in explaining subjects’ behavior. The superiority of the Late-Inference model was further supported by a model comparison using a Bayesian information criterion (BIC; **Supplementary Table 2**).

**Figure 4.**
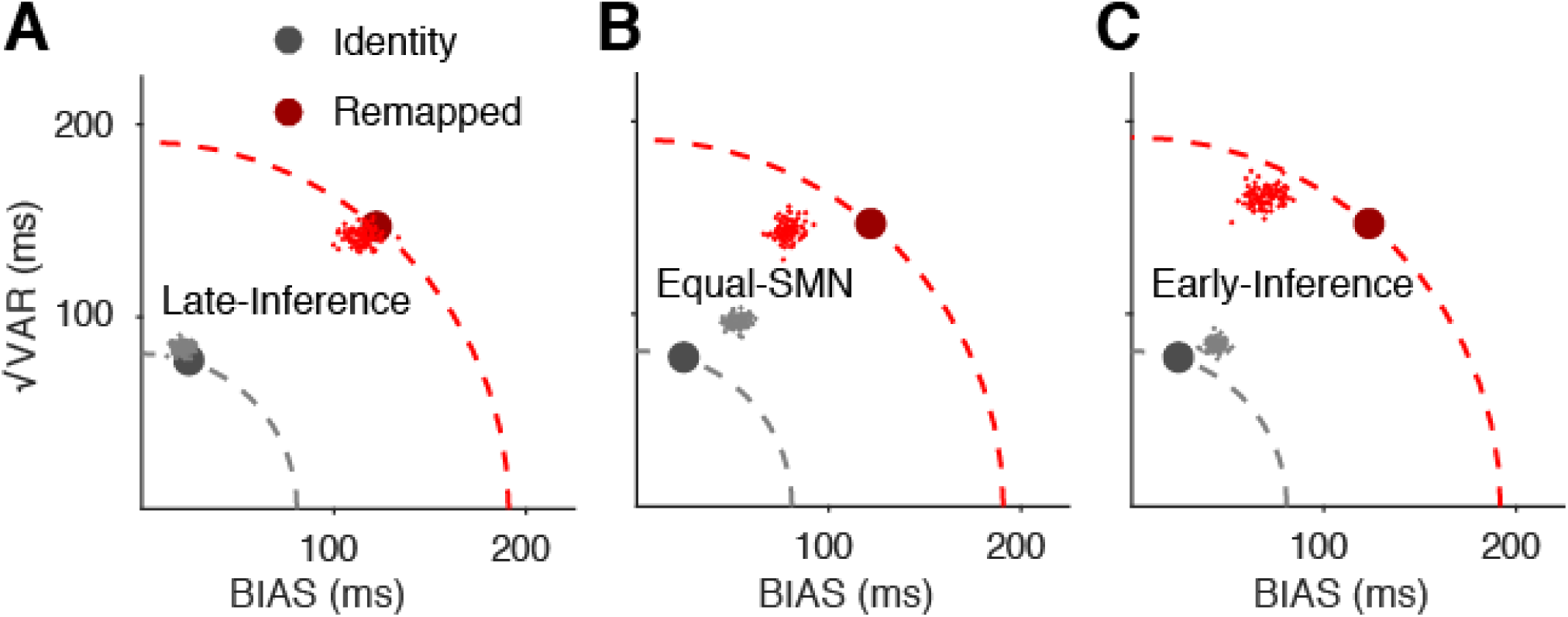
Model comparison for the time measurement and production task. (**A**) Comparison of *BIAS* and *√VAR* of a typical subject to simulations generated from fits to behavior using the Late-Inference model. The model uses the same measurement and production Weber parameters (*w*_*m*_ and *w*_*p*_) across the identity (gray) and remapped contexts (red) and introduces additional scalar SMN in the remapped context parameterized by *w*_*t*_. Large solid circles represent the subject’s behavior and small dots represent individual simulations (n = 100). (**B**) Same as **A** for the Equal-SMN model. This model was identical to the previously described Bayesian observer model (**Supplementary Figure 1; Figure 3**) with *w*_*m*_ and *w*_*p*_ but no *w*_*t*_. The failure of this model to capture subjects’ behavior rejects the null hypothesis that the remapped context can be explained without additional SMN. (**C**) Same as **A** for the Early-Inference model. This model includes *w*_*m*_, *w*_*p*_, and *w*_*t*_, but the additional scalar SMN is introduced after the inference stage. The Early-Inference model failed to capture the disparity in *BIAS* between the two contexts when compared with the Late-Inference model, further supporting the late inference hypothesis.

We considered a number of other models, but were unable to create any model that could account for the increased BIAS as accurately as the Late-Inference model. Here, we describe two alternative models that predicted some additional bias in the presence of higher SMN but were nonetheless inferior to the Late-Inference model. The first model, which we refer to as “Late-Ignore-SMN”, is a variant of the Late-Inference model in which the observers makes the inference after the addition of SMN (through *w*_*t*_), but does not take SMN into account. In other words, this is a model of an observer that uses a Late-Inference strategy but does not have an internal model of SMN. The second, which is referred to as the “Observer-Actor” model (Jazayeri and Shadlen 2010; Acerbi, Wolpert, and Vijayakumar 2012), is a variant of the Early-Inference model, which is additionally optimized for scalar variability in the production stages, but not for SMN. Both of these models predicted some degree of increased bias and a substantial (and suboptimal) increase in variability that failed to capture subjects’ behavior in the remapped context (**Supplemental Figure 3, Supplementary Table 2**).

Another explanation that might account for the increased bias in the remapped context is that subjects did not learn the transformation correctly, and instead of applying a gain, simply added a fixed delay to their responses irrespective of the sample interval. Such an offset-adjustment strategy would result in an effective increase in bias and could thus masquerade as a late Bayesian inference strategy. To investigate this possibility, we designed a control experiment with a gain factor of 0.75 instead of 1.5. For a gain of 0.75, a similar offset-adjustment strategy would require subjects to subtract a fixed delay from their responses. This would predict that responses would exhibit less bias than predicted from scaling responses by a factor of 0.75 (i.e., the prediction from the equal SMN hypothesis). However, for the gain of 0.75, similar to the case for a gain of 1.5, subjects’ *RMSE*, *BIAS*, and fits to *w*_*m*_ were higher than predicted by the identity context (**Supplementary Figure 4**). Therefore, the increased bias could not be explained by an offset-adjustment strategy.

## Length measurement and production task

In a second experiment, we asked whether SMN degrades performance in a length production task, and whether humans use a late inference strategy to optimize performance in the presence of SMN. To answer this question, we tested the behavior of seven subjects in a task that involved drawing a line whose length was either matched to (identity context) or 1.5 times (remapped context) the length of a visually presented bar (**Figure 5A**). The length of the bar was sampled from a discrete uniform distribution with 11 values ranging between 10 and 15 visual degrees and was presented on a horizontally oriented monitor. After presentation of the visual bar, subjects had to draw a bar by moving a handle that controlled the position of a cursor on the monitor (**Figure 5A**).

**Figure 5.**
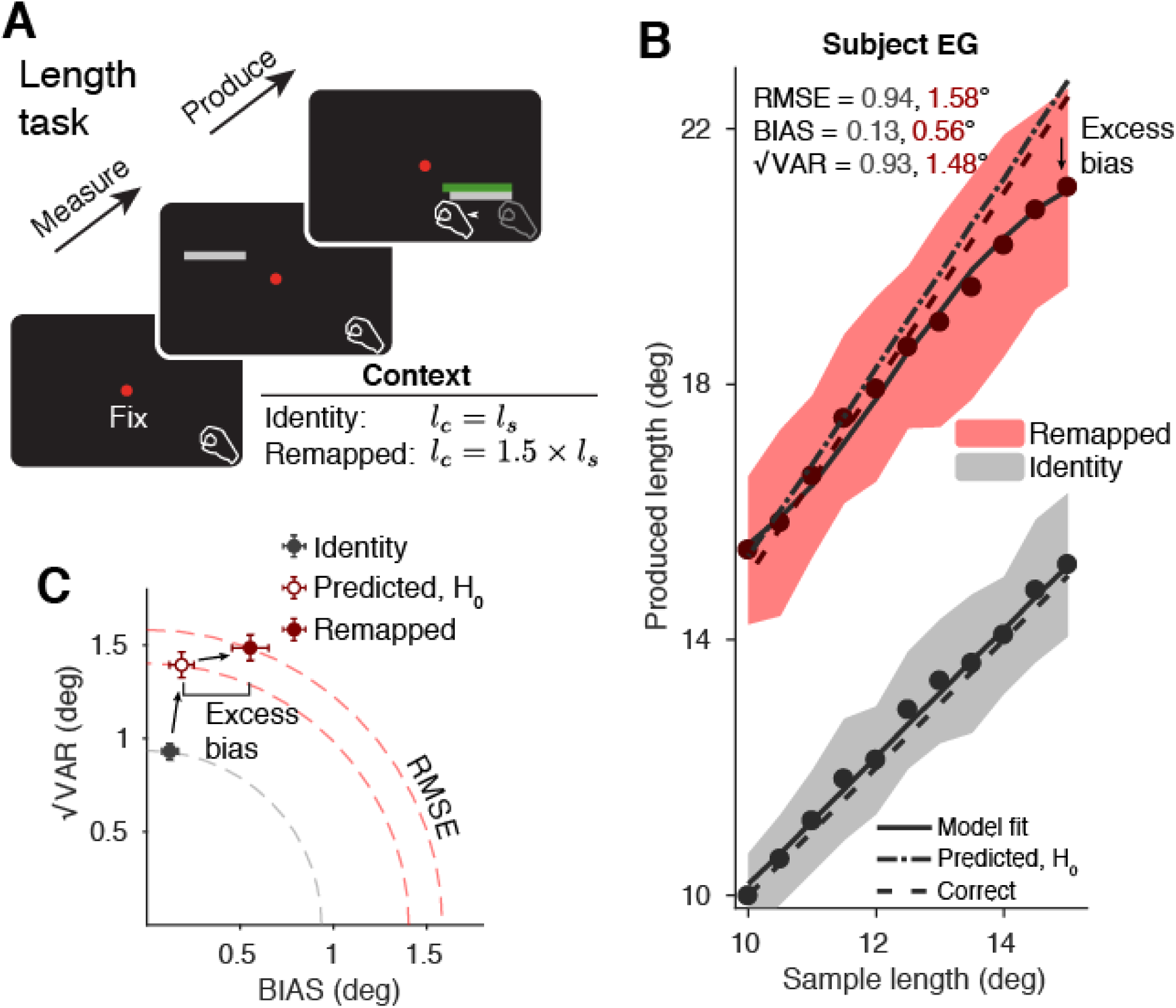
Length measurement and production task. **A**. Trial structure. Each trial began with the presentation of a red fixation spot. Subjects first measured the length *l*_*s*_ of a gray sample bar presented briefly on an upwards facing monitor. After the sample bar was extinguished, subjects moved a manipulandum containing a digitizing pen located under the monitor in order to draw a bar which was as close in length as possible to the correct length *l*_*c*_ = *g × l*_*s*_. As in the timing task, there were two contexts: in the identity context, *gain* = 1, whereas in the remapped context, *gain* = 1.5. After the response, subjects received scaled and signed feedback via the presentation of a gray bar of the proper length while the subjects’ produced bar changed color to red (for an inaccurate response) or green (for an accurate response; see Methods). **B**. Performance of an example subject in the identity (gray) and remapped (red) contexts (same format as **Figure 2**). Filled circles and shaded regions indicate mean response times ± one standard deviation; dashed lines represent correct intervals. Solid lines represent the mean responses of a Bayesian observer model (see Methods) fit to the subject’s data separately for the two contexts; the dash-dot line in the *gain* = 1.5 condition corresponds to the prediction for the remapped context using parameters of the model fit to the identity context (H_0_: no additional SMN). Comparing the model fit for the remapped context to the no additional SMN prediction, the excess bias towards the mean in the remapped relative to the identity context can be seen. **C.** *√VAR* vs. *BIAS* for the two contexts (gray for identity, and dark red for remapped), as well as the prediction for the remapped context assuming no additional SMN (empty circle). This prediction underestimates *RMSE* indicating larger SMN for the remapped context. The increased RMSE was predominantly due to an increase in BIAS (“excess bias”). Dashed quarter circles illustrate combinations of *BIAS* vs. *√VAR* giving rise to equal *RMSE*; error bars are computed from the standard deviation of bootstrapped estimates (n = 1000).

**Figure 5B** shows the behavior of one subject in the length measurement and production task. Similar to the timing task, *RMSE* was higher in the remapped context compared to the identity context suggesting that SMN was larger for the transformation associated with *gain* = 1.5. Moreover, the increase in *RMSE* was associated with an excess bias beyond what was expected from multiplying the observed bias in the identity context by the gain. The higher *RMSE* and *BIAS* was a general finding across subjects (**Figure 6A**) indicating that the length task was also associated with a late Bayesian strategy to compensate for the larger SMN.

**Figure 6.**
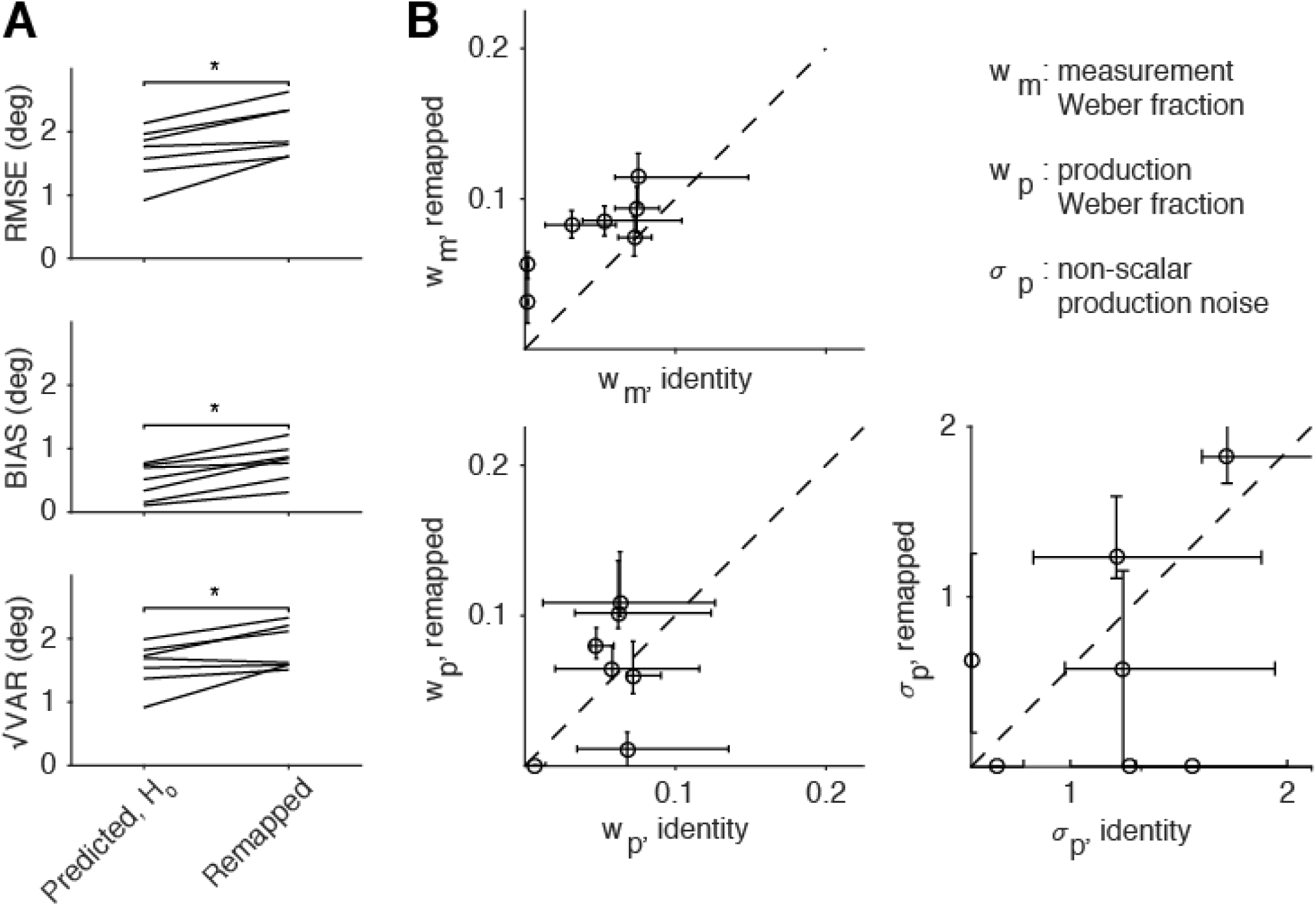
Summary of subjects’ performance in the length production task. (**A**) Comparison of *RMSE*, *BIAS*, and *√VAR* for all subjects. The lines connect values predicted in the remapped context based on simulated data from the observer model fit to individual subjects’ behavior for gain of 1 and assuming no additional SMN (predicted, H_0_; left) to actual values observed from behavior (*gain* = 1.5; right). Similar to the timing task, subjects had higher error and were more biased towards the mean than was predicted assuming no additional SMN in the remapped context. Variability also increased in most subjects (*: p < 0.05). (**B**) Fitting the observer model independently across the two contexts resulted in higher values of *w*_*m*_ in the remapped context, reflecting additional reliance on prior information consistent with a late inference strategy. Values for *w*_*p*_ and the non-scalar motor noise parameter σ*p* were not systematically affected by the gain. Error bars represent 95% confidence intervals estimated using a bootstrap procedure (n = 1000).

To further validate these results using model comparisons, we first sought to develop an ideal observer model of the length task. In the timing task, the Bayesian observer model we used was based on previous work using a time reproduction task (Jazayeri and Shadlen 2010) and included two parameters: one scaling factor for the measurement noise (*w*_*m*_), and another for the production noise (*w*_*p*_). For the length task, we considered the possibility that the production stage might be subject to additional execution noise due to hand movements, as previous work has suggested (Wolpert, Ghahramani, and Jordan 1995; Robert J. van Beers, Haggard, and Wolpert 2004). We compared a model similar to the timing task with scalar measurement and production noise to another model that included an additional signal independent production noise term σ*p*. The model with the additional nonscalar production noise provided a better description of behavior despite having an additional parameter (relative BIC of 64, compared to 49 in favor of pure scalar model for the timing task). We therefore proceeded with this scalar-nonscalar model to compare the identity and remapped contexts.

Similar to the timing task, observer model fits in the remapped context were associated with higher values for *w*_*m*_ and no systematic relationship to *w*_*p*_ and σ*p* (**Figure 6B**). These results indicate that the higher SMN in the remapped context is accompanied by higher reliance on prior information, as expected from the late Bayesian inference strategy. This result was further substantiated by a direct model comparison based on BIC showing that the Late-Inference model was best at capturing the data in the remapped context (see **Supplementary Tables 3 & 4** for individual subjects’ results and summary).

## Discussion

Noise in sensorimotor transformations directly impacts performance. To optimize behavior in the presence of sensorimotor noise, the brain must adopt a late inference strategy that takes sensorimotor noise into account and adjusts motor plans according to the statistics of the outcomes. Our results indicate that subjects compensate for noise in sensorimotor transformations by biasing responses towards the mean of a sensorimotor prior, supporting the hypothesis that humans seek to optimize behavior in the presence of sensorimotor noise by adopting a late inference strategy. This finding extends previous work on Bayesian models of sensory and motor systems, and indicates that the brain circuits are additionally optimized for noise arising from sensorimotor transformations.

The vast majority of experiments on the application of Bayesian theory to behavior involve some sort of sensorimotor transformation. As we found in our work, and others found in other behavioral settings (Soechting and Flanders 1989b; Gordon, Ghilardi, and Ghez 1994; Pine et al. 1996; McIntyre et al. 2000; Sober and Sabes 2005; Churchland, Afshar, and Shenoy 2006; Schlicht and Schrater 2007), sensorimotor transformations are often noisy. Surprisingly however, most Bayesian models do not take SMN into account and formulate Bayesian calculations in terms of sensory inference (Weiss, Simoncelli, and Adelson 2002; Tassinari, Hudson, and Landy 2006; Jazayeri and Shadlen 2010; Ganguli and Simoncelli 2014). This approach may be adequate when the sensory noise is the dominant source of uncertainty (Osborne, Lisberger, and Bialek 2005). However, sensorimotor transformations may generate a substantial fraction of the total noise. Indeed, for the Late-Inference model, which captured behavior most accurately, the transformation noise (*w*_*t*_) associated with the remapped context was larger than measurement noise (*w*_*m*_; **Supplementary Tables 2 & 4**). This suggests that many previous experiments that ignored SMN and yet found the behavior to be optimal might have misattributed SMN to noise in sensory representations (see **Figures 3B & 6B**). This is not surprising as distinguishing between sensory and sensorimotor noise is not straightforward when the two are not independently manipulated. Our experiment was designed to overcome this challenge by comparing identity and remapped sensorimotor contexts and thus manipulating SMN without changing the sensory noise. Finally, it is important to note that our own work is not fully immune to the misattribution of SMN to sensory noise. While we were able to reveal the excess bias due to larger SMN in the remapped context, it is conceivable that SMN was a significant factor in the identity context as well. This might explain why a previous study found time measurement and reproduction task to be more biased than predicted by noise levels in a temporal bisection task (Cicchini et al. 2012). As such, we might have underestimated the importance of SMN by misattributing some portion to SMN to measurement noise in both contexts.

Our proposal of a late inference strategy unifies various observations in a wide range of sensorimotor tasks. For example, it has been shown that when both visual and proprioceptive information are available, subjects rely more strongly on the modality that had the least transformational complexity (Sober and Sabes 2005). Schlicht and Schrater (Schlicht and Schrater 2007) showed that subjects account for the effects of eye position uncertainty in a grasping task by increasing grip aperture. Another study found that reach movements in 3-dimensional space were consistently biased towards the centroid of target distributions, particularly along the radial (distance) axis, and that this bias was not seen when subjects performed a simpler pointing task which only required wrist movements (Soechting and Flanders 1989b). The authors’ interpretation of these results was that the brain implemented linear approximations to the true nonlinear transformations between target location and motor commands (Soechting and Flanders 1989a). However, our results suggest that the bias might be due to a stronger influence of prior information when facing the more challenging sensorimotor task of reaching in 3D. Late inference may also explain response patterns in reaching tasks where biases exist in arm-centered, rather than eye-centered reference frames (Baud-Bovy and Viviani 1998), or in cases where target-dependent bias unexplained by sensory noise is attributed to suboptimal aiming strategies (Tassinari, Hudson, and Landy 2006).

Moreover, the late inference model provides a natural explanation for why various post-sensory cognitive operations cause additional biases in behavior. Without a late inference strategy, one would expect sources of noise such as memory decay to lead to additional variability. However, a number of experiments have shown that post-sensory noisy computations can lead substantial increases in bias. Examples include mental operations in the presence of memory delays (Moyer et al. 1978; Ashourian and Loewenstein 2011), predictions of complex kinematics (Smith and Vul 2013; Battaglia, Hamrick, and Tenenbaum 2013), and pointing in 3D space under various memory loads (McIntyre et al. 2000). In all these cases, the additional biases can be straightforwardly accounted for by considering the possibility that the brain has an internal model of post-sensory noisy transformations and compensates for them by using its prior knowledge about the desired outcomes.

Late inference may seem at odds with recent proposals that priors based of natural stimuli may already be applied at low-level sensory areas (Weiss, Simoncelli, and Adelson 2002; Girshick, Landy, and Simoncelli 2011). However, the late inference model is fully compatible with early integration of sensory inputs with priors as well as multiple stage of updating the posterior (Ma et al. 2006; Beck et al. 2008; Ganguli and Simoncelli 2014; Ma and Jazayeri 2014). The key constraint imposed by the late inference strategy is simply for the brain to delay extracting a point estimate from the posterior distribution to as late as possible (Simoncelli 2009).

Finally, our results bear on the computational principles that govern brain function when information undergoes multiple processing stages. We found that the brain computations are optimized for sensorimotor transformations suggesting that the inference is made after the addition of sensorimotor noise. However, late inference is not a special requirement that only applies to sensorimotor noise. The passage of information in the simplest visuomotor task from the primary visual cortex to downstream visual areas to sensorimotor cortex to movement control circuits undergoes many stages of processing. Each stage of processing is likely to have its own private noise and can thus add to the overall variability. Regardless of the task and where the noise is added, the optimal strategy is to delay inference until after the final stages of processing (Simoncelli 2009). This applies even when intermediate transformations are trivial, as must be the case when most of the error can be attributed to sensory processing (Osborne, Lisberger, and Bialek 2005). This optimality consideration coupled with evidence from our work that the brain does indeed delay inferences until after the introduction of sensorimotor noise suggest that brain circuits may employ Bayesian inference as an inherent computational principle.

## Methods

### Subjects

Human subjects aged 18-65 years participated in this study after giving informed consent. All experiments were approved by the Committee on the Use of Humans as Experimental Subjects at the Massachusetts Institute of Technology. The study consisted of two experiments: a time interval estimation and production task (Experiment 1), and a length estimation and production task (Experiment 2). A group of 9 subjects participated in Experiment 1 (8 for gain = 1.5 and 9 for gain = 0.75), and a mostly different group of 7 subjects participated in Experiment 2 (1 subject participated in both experiments). All subjects had normal or corrected-to-normal vision.

### Procedures

Experimental sessions lasted approximately 45-60 minutes. Each subject completed 1-2 sessions per week. Experiments were controlled by an open-source software (MWorks; http://mworks-project.org/). All stimuli were presented on a black background. Although eye movements were not monitored, all trials began with central fixation spot that subjects were asked to hold their gaze on throughout the trial. In Experiment 1, subjects viewed all stimuli binocularly from a distance of approximately 67 cm on either a 23-inch Apple A1082 LCD monitor at a resolution of 1900 × 1200 driven by an Intel Macintosh Mac Pro computer, or a 24-inch early 2009 Apple Mac Pro at a refresh rate of 60 Hz in a dark, quiet room. In this experiment, responses were registered on a standard Apple Keyboard connected to the experimental machine. In Experiment 2, subjects viewed stimuli from above on a 21.5-inch Samsung SyncMaster SA200 monitor, and responses were registered using a pen digitizer tablet (Wacom Intuos5 touch); the stylus was fixed at a vertical position inside a custom printed handle which subjects grasped.

### Behavioral tasks

Our objective in both experiments was to investigate the effect of sensorimotor noise (SMN) on performance. To do so, each experiment consistent of two sensorimotor contexts, an “identity” context, and a more challenging “remapped” context that was expected to involve higher levels of SMN. In each experiment, human subjects measured a scalar sensory quantity (time interval in Experiment 1, and length in Experiment 2) drawn from a prior distribution. In the “identity” context, subjects had to reproduce the sensory quantity (the sample), and in the “remapped” context, they had to produce the same quantity multiplied by a gain factor. Subjects whose responses for the shortest and longest stimuli in the identity context were at least one d’ (d-prime) apart were invited to participate in the main experiment.

#### Experiment 1: Time interval estimation and production task

The behavioral task used in Experiment 1 was a variant of the Ready, Set, Go task used in a previous study (Jazayeri and Shadlen 2010). Subjects had to measure a sample interval drawn from an 11-point discrete uniform distribution between 600 and 1000 ms, then immediately produce an interval that was equal to the sample interval multiplied by a gain factor. The sample interval was demarcated by two visual flashes (“Ready” and “Set”) located to the left and above a fixation point at the center of a computer monitor. The production interval was defined as the interval between the onset of the second flash and the response (key press) of the subject. In the identity context, the gain was 1, whereas in the remapped contexts the gain was either 1.5 or 0.75. The gain was fixed in each behavioral session and was communicated at the beginning of each session as either “same,” “shorter,” or “longer.” The gain was also evident on every trials: the ratio of the horizontal distance between the Ready flash to the left of the fixation point and a “Go” cue to the right fixation was set by the gain factor. Following each response, subjects were given feedback regarding their response via a round marker displayed a distance from the Go cue proportional to the error and regarding the trial outcome via the color of the marker. A green marker indicated a “hit” and a white marker indicated a “miss.” Subjects completed two consecutive sessions of 600 trials for each gain; the hit/miss threshold was on a staircase for the first session and fixed for the second session at the mean of the last 100 trials of the first session. Analyses were performed using data from the second sessions. All subjects completed the “identity” context sessions first, followed by either the gain of 1.5 or gain of 0.75, selected pseudorandomly for each subject.

#### Experiment 2: Length estimation and production task

The behavioral task used in Experiment 2 was conceptually similar to the first in that subjects produced a scalar quantity multiplied by a gain factor. However, instead of a time interval, subjects measured and produced visually presented lines drawn from an 11-point discrete uniform distribution between 10 and 15 degrees visual angle. To produce the length, subjects had to move a manipulandum underneath a horizontally positioned computer monitor. In each trial, after subjects positioned the manipulandum at the perimeter of the screen, a horizontal line flashed for 500 ms, after which subjects had 1200 ms to move the manipulandum inward to the final response position. Two small vertical bars, one positioned at the initial location of the manipulandum and one tracking the horizontal location of the bar, provided online visual feedback during the response. The produced length was measured as the distance between the two vertical bars at the end of the response period. The gain in the identity and remapped contexts was 1 and 1.5, respectively. The gain was communicated by telling subjects to produce either “the same as” or “one and a half times” the length of the sample. Response feedback was similar to the interval production task: following each response, the produced length was shown as a line between the marker bars (green for hit and red for miss), and the correct length was displayed immediately beneath in gray. Subjects completed four sessions total with each session comprising two blocks of 150 trials of identity and remapped trials for a total of 600 trials per session. The error threshold for each gain was on a one-up one-down staircase for the first two sessions and fixed for the final two sessions at the mean of the last 100 trials for each gain. The order of blocks associated with the identity and remapped blocks was pseudorandomized across subjects.

### Data analysis

Behavioral performance in all tasks was quantified with three statistics (Jazayeri and Shadlen 2010): *BIAS*, *√VAR*, and *RMSE*. *BIAS* summarizes the difference between average and correct responses and is defined as

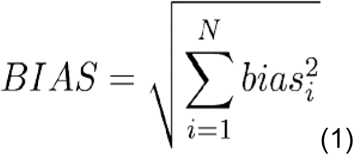

where *bias* is the difference between the mean response and correct response for a given sample interval. *√VAR* summarizes the variability of responses:

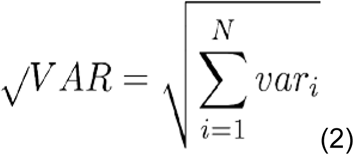

where *var* is the variance of the responses for a particular sample interval. Because samples were drawn randomly, it was not the case that the number of trials for each sample was exactly the same. Therefore, averages of for *BIAS* and *√VAR* were normalized across samples according to the number of trials presented. Finally, *RMSE* was calculated as:

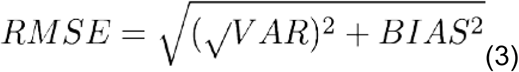

The three quantities are related through a sum of squares: *RMSE*^2^ *=* (*√VAR*)^2^ *+ BIAS*^2^ (see **Figure 1B**). Prior to analyzing data, we identified and removed “lapse” trials for each subject. This involved finding and removing trials for which responses were greater than three standard deviations from the mean response for a particular sample quantity and context, and which was performed twice iteratively.

### Model descriptions and fitting procedure

We employed a Bayesian model previously shown to capture the behavior of human subjects in the timing task (Jazayeri and Shadlen 2010). The model consists of three stages: a noisy measurement stage, a deterministic Bayesian inference stage, and a noisy production stage (**Supplementary Figure 1**). The noisy measurement *t*_*m*_ (*t* = time) is generated according to the noise model p(*t*_*m*_|*t*_*s*_), then used to generate an inference *t*_*i*_ which minimizes the expected squared error between *t*_*i*_ and *t*_*s*_ given *t*_*m*_:

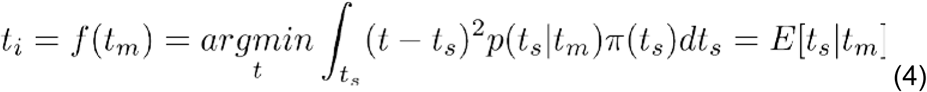

Where π(t_s_) represents the observer’s “prior” belief about *t*_*s*_. The inference, which can be thought of as a perceptual estimate, is the expected value of the sample interval given the measurement. The model then generates *t*_*p*_ according to the production noise model p(*t*_*p*_|*t*_*i*_). p(*t*_*m*_|*t*_*s*_) and p(*t*_*p*_|*t*_*i*_) were formulated as Gaussian distributions with means *t*_*s*_ and *t*_*i*_, respectively, and standard deviations that scaled with the respective means. This model has two free parameters, *w*m and *w*_*p*_, which represent the Weber fractions (i.e., ratio of standard deviation to mean) for p(*t*_*m*_|*t*_*s*_) and p(*t*_*p*_|*t*_*i*_), respectively. Generally, the model captures high response variability by increases in *w*_*p*_, and large response biases towards the mean of the prior by increases in *w*_m_. As a corollary, increases in *w*_*p*_ have a comparably larger effect on total error (**Supplementary Figure 2**).

This model was used in three ways. First, we used it to predict responses in the remapped context from fits of the model to the identity context (without changing the model parameters). We used this approach to generate predictions for the null hypothesis that the remapped context did not engender additional SMN. Second, we fit this model to the behavior but allowed *w*_*m*_ and *w*_*p*_ to take on different values in the two contexts. We used this approach to distinguish between the early and late inference hypotheses. Based on the behavior of this model (**Supplementary Figure 2**) we expected a late inference strategy would cause an increase in response biases and lead to systematic increases in the fit to *w*_*m*_ but not *w*_*p*_. Third, we used the model to fit the data combined across the two contexts. This approach would succeed under the null hypothesis that the two contexts have the same level of SMN. We refer to this model the “Equal-SMN” model.

We also developed an “Early-Inference” and a “Late-Inference” model, which included an additional parameter, *w*_*t*_, to capture putative additional SMN in the remapped context. These models were fitted to the combined data in the two contexts. In the Early-Inference model, Bayesian inference was applied to the sensory measurement stage, and SMN was added to the production stage (after inference). To do so, we formulated the production stage such that *p(t*_*p*_*|t*_*i*_) was a scalar Gaussian distribution with mean of *gain × t*_*i*_ and modified Weber fraction 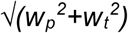. In contrast, for the Late-Inference model, the effect of SMN was applied prior to the inference stage by drawing samples of the “remapped” measurement from a Gaussian distribution *p(t*_*t*_*|t*_*s*_) with mean of *gain × t*_*s*_ and Weber fraction 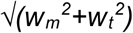, where *t*_*t*_ (transformed interval) represents *t*_*m*_ multiplied by the remapping gain.

In the Early-Inference model, the prior distribution corresponded to the sensory variable (i.e., prior to transformation), whereas in the Late-Inference mode it was formulated based on the transformed sensory variable (i.e., sensory variable multiplied by the gain). This modified prior represents the observer’s belief about the correct interval and can be viewed as a sensorimotor rather than sensory prior.

Model parameters were fit by maximizing the log-likelihood of subjects’ responses given the sample values and gain. The maximization was done using the fminsearch command in MATLAB (Mathworks). Model fitting and simulation involved numerical integration over the posterior distribution using Simpson’s rule. Parameter searches were repeated ten times each with different parameter initialization, and results were inspected for consistency.

## Acknowledgements

We would like to thank Josh McDermott, Seth Egger, and Devika Narain for valuable comments regarding the manuscript. M.J. is supported by NIH (NINDS-NS078127), the Sloan Foundation, the Klingenstein Foundation, the Simons Foundation, the Center for Sensorimotor Neural Engineering, and the McGovern Institute.

## Manuscript Supplement

### Additional models

We considered two additional models: the Late-Ignore-SMN model and the Observer-Actor model, both of which predicted more bias than the Early-Inference and Equal-SMN models. However both models also predicted a substantial increase in variance (**Supplemental Figure 3**), which deviated from subjects’ behavior.

### The Late-Ignore-SMN model

This model utilizes a late inference stage, but the additional noise from sensorimotor transformations is ignored. More concretely, the model assumes that variability prior to inference is determined by an effective Weber fraction of 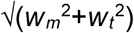 but that subjects compute the posterior based on the incorrect assumption that the Weber fraction was *w*_*m*_. In other words, the inference function *f*() ignores *w*_*t*_. The reason why ignored SMN causes an increase in bias in the Late-Ignore model may be somewhat counterintuitive. To understand this, let us examine the computations that underlie Bayesian inference in the presence and absence of SMN. In the absence of SMN, there would be a one-to-one correspondence between a measurement and its transformation. For measurements (and transformations thereof) that are farther away from the mean of the uniform prior, the nonlinear inference function (**Supplementary Figure 1, middle panel**) causes more bias in the inferred values. However, the magnitude of this bias would be the same for both early and late inference because of the one-to-one correspondence between a measurement and its transformation. This situation changes however in the presence of SMN. SMN makes the result of this transformation stochastic such that a single measured interval leads to a distribution of transformed measurements. Therefore, the magnitude of bias associated with a measurement has to be computed as an expectation across the distribution of transformed measurements. Since the inference function is nonlinear, it disproportionately biases transformed measurements that are farther away from the mean of the prior distribution (and in particular, outside the support of the uniform prior) leading to an overall increase in the magnitude of the bias.

While the Late-Ignore-SMN model was able to accommodate more bias, it was not enough to capture the excess bias observed in the data. The model also predicted substantially increased variance, as well as a skewing of response distributions away from the correct response (see **Supplementary Figure 3**). We also examined the behavior of the Late-Ignore-SMN model under the assumption that observers approximate uniform priors as Gaussian-like functions as was suggested previously (Acerbi, Wolpert, and Vijayakumar 2012; Cicchini et al. 2012). This possibility reduces the amount of bias introduced by the Late-Ignore-SMN model (data not shown), and further diminishes its utility as a viable model for the observed biases in subjects’ behavior.

### The Observer-Actor model

This is a variant of the Early-Inference model for which inference is made after the sensory measurement stage and the production stage is modeled by a Gaussian with mean of *gain × t*_*i*_ and Weber Fraction 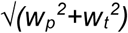. However, following previous work showing that humans account for response variability when planning actions (Trommershäuser et al. 2005; Landy, Trommershäuser, and Daw 2012) we augmented the inference stage so that the early inference would take into account the scalar post-inference variability due to the transformation and production (Acerbi, Wolpert, and Vijayakumar 2012). The optimal inference under the Observer-Actor model can be computed by marginalizing over the distribution of produced values, *p(t*_*p*_*|t*_*i*_):

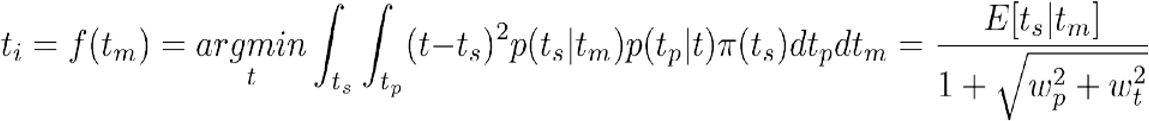

Here *t*_*i*_ is the optimal value that the observer using early inference should aim to produce in order to mitigate the effects of post-inference transformation and production scalar variability. Like the Late-Ignore-SMN model, this model was able to accommodate a slight increase in bias as it specified all aimed magnitudes to be smaller than dictated by the Early-Inference model. However, also similar to the Late-Ignore-SMN model, increasing SMN (*w*_*t*_) primarily affected variance rather than bias (**Supplemental Figure 3**).

**Supplementary Figure 1.**
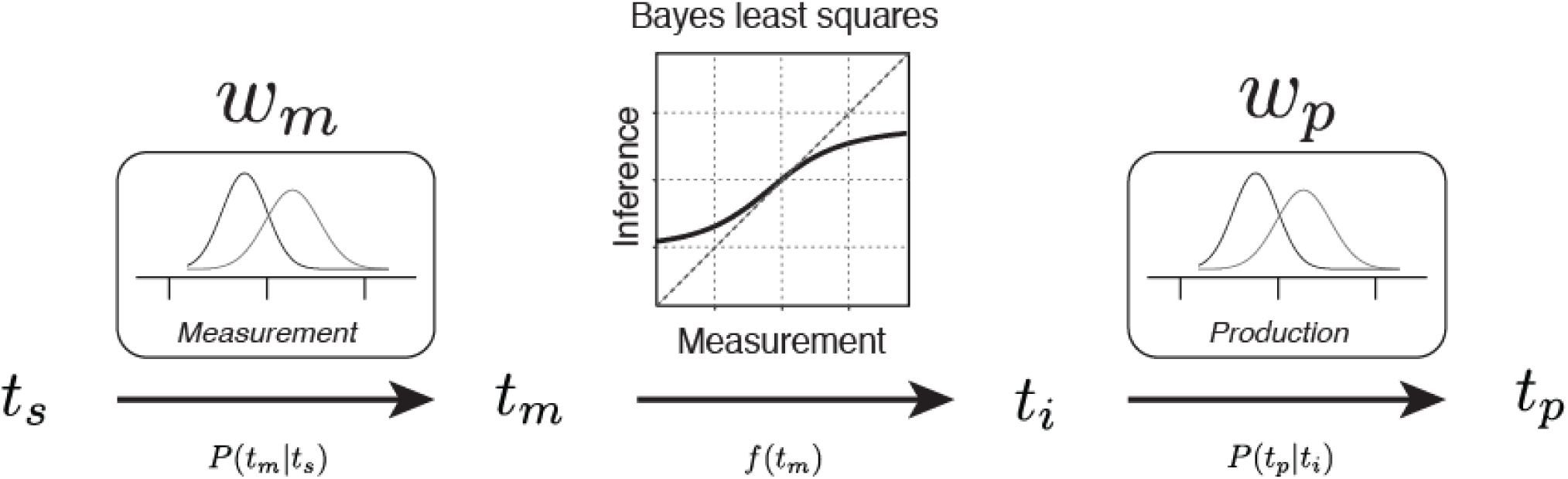
Illustration of the Bayesian observer model previously shown to capture the behavior of human subjects in the identity context of the timing task (Jazayeri and Shadlen 2010). The noisy measurement stage (left), takes a sample interval (*t*_*s*_) as input and produces a measurement (*t*_*m*_) corrupted by Gaussian noise with standard deviation equal to the measurement Weber fraction (*w*_*m*_) times the value of *t*_*s*_. A deterministic Bayesian inference stage (middle) then takes *t*_*m*_ as input and produces an inference *t*_*i*_ using knowledge of the prior distribution over *t*_*s*_ as well as the value of *w*_*m*_ such that the the root mean squared error (RMSE) of *t*_*i*_ relative to *t*_*s*_ is minimized. Finally the noisy production stage (right) takes *t*_*i*_ as input and generates a produced time (*t*_*p*_) corrupted by Gaussian noise with standard deviation equal to the production Weber fraction (*w*_*p*_) times the value of *t*_*i*_. This model captures high response variability by increases in *w*_*p*_ and large response biases towards the mean of the prior by increases in *w*_m_ (**Supplementary Figure 2**).

**Supplementary Figure 2.**
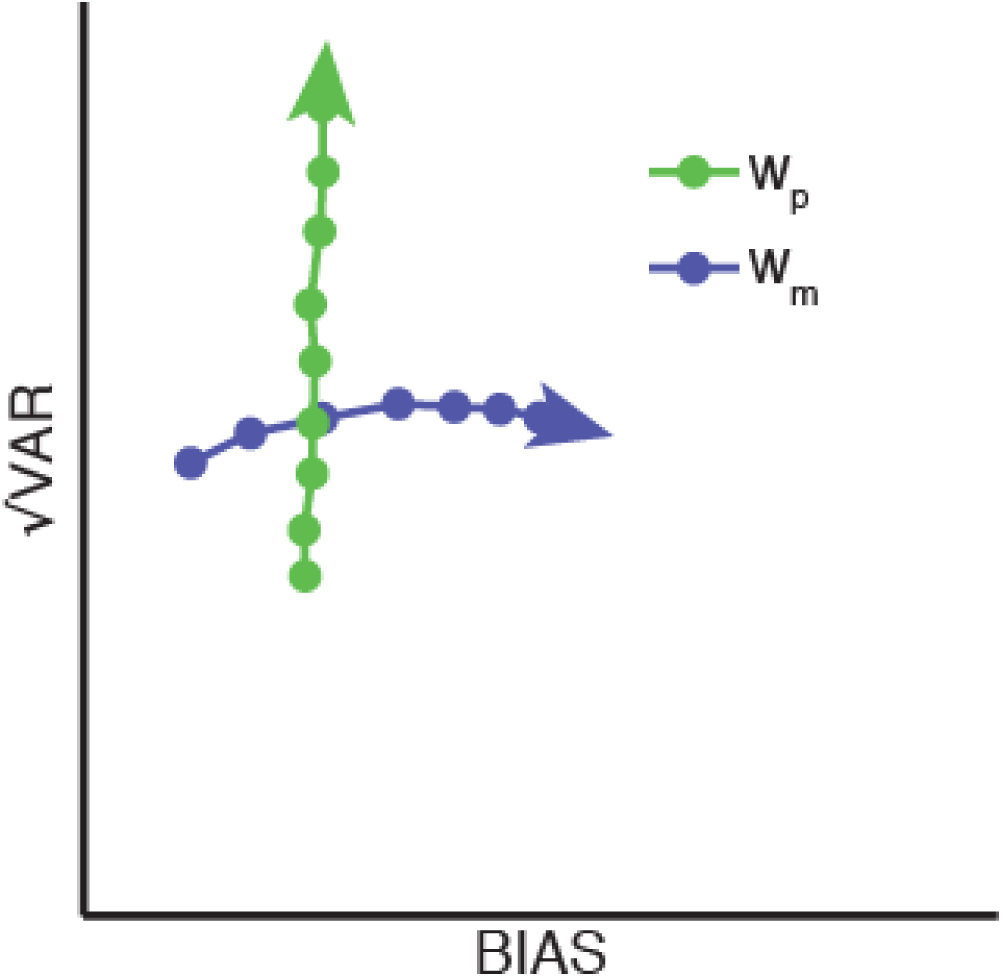
Illustration of the effects of increasing the amount of production and measurement noise. Using the Bayesian observer model (Jazayeri and Shadlen 2010), we simulated the effects of varying the amount of motor production noise (green; *w*_*p*_: production Weber fraction) and sensory measurement noise (blue: *w*_*m*_: measurement Weber fraction) on *√VAR* and *BIAS*. Increasing the value of *w*_*p*_ substantially increased variability with almost no increase in bias, whereas increasing *w*_*m*_ increased bias due to increased reliance on prior information, but had little effect on variability. Thus, we interpret increases in *w*_*m*_ (**Figures 3 & 6**) for subjects in the remapped contexts as evidence for reliance on prior information in a late inference strategy to mitigate the effects of increased SMN relative to the identity contexts.

**Supplementary Figure 3.**
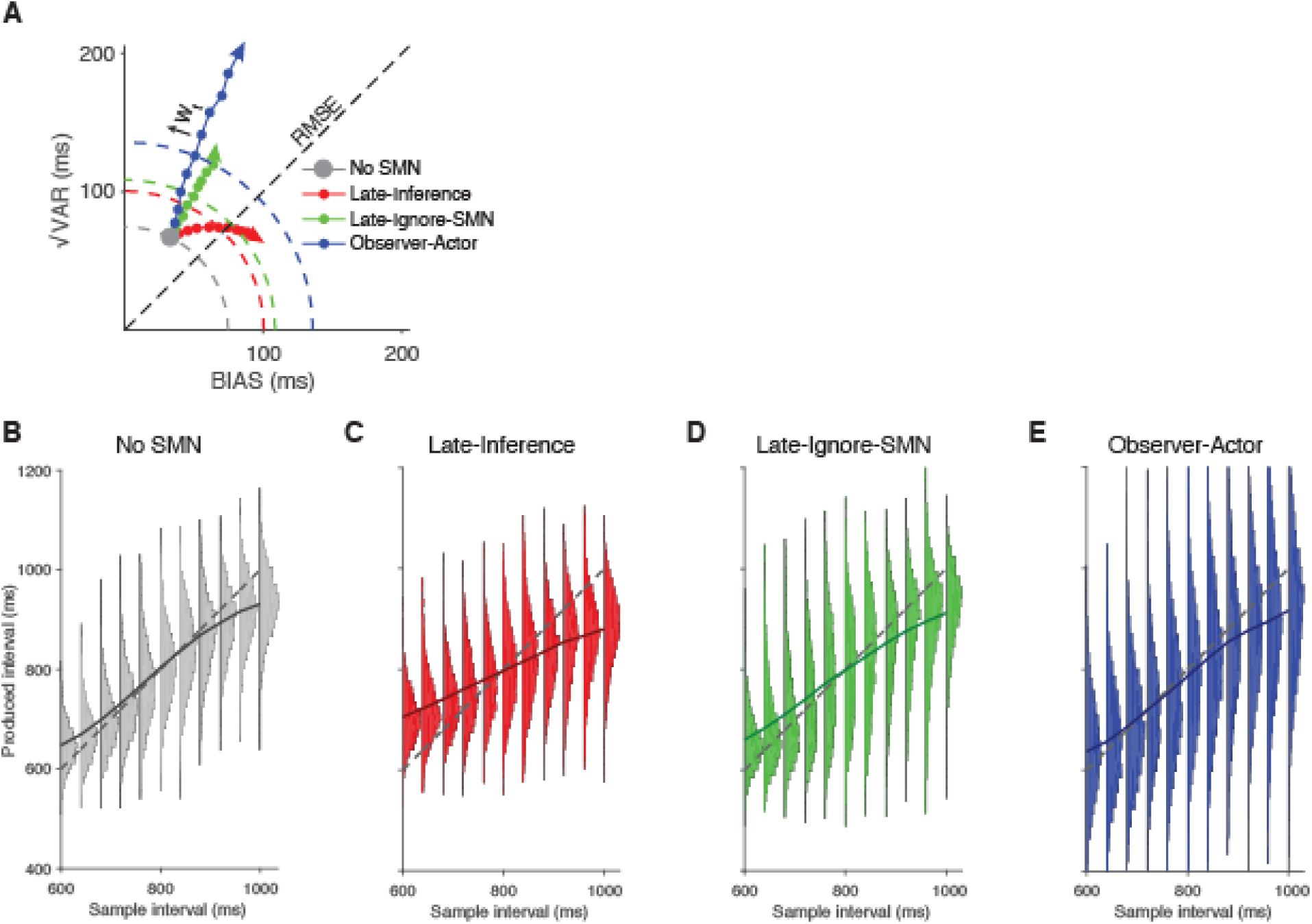
Behavior of different models as a result of increasing SMN. **A**. Bias variance curves for three models with increasing SMN (*w*_*t*_ *=* 0.025 - 0.25) relative to a “no SMN” observer model (large gray circle; *w*_*m*_ = 0.08; *w*_*p*_ = 0.06). For each model (Late-Inference, red; Late-Ignore-SMN, green; Observer-Actor, blue) increasing SMN (*w*_*t*_) causes *RMSE* to increase as evident by increasing distance of circles from the origin. The Late-Inference model compensates for increasing SMN by systematic increases in *BIAS* and has the best performance. The other models exhibit less BIAS and more *√VAR* leading to larger *RMSE* compared to the Late-Inference model. Quarter circles represent combinations of *√VAR* and *BIAS* with equal *RMSE* for *w*_*t*_ = 0.14. **B-D.** Distribution of responses as a function of sample duration under various models for *w*_*t*_ = 0.14. **B.** The no SMN observer model, which shows the lowest *RMSE*. **C.** The Late-Inference model (red). In this model, bias increases substantially while variability remains relatively stable (similar to the effect of increasing *w*_*m*_; **Supplementary Figure 2**). **D.** The Late-Ignore-SMN model also adds bias; however, it also leads to a comparable increase in variance. **E.** Increasing transformation noise in the Observer-Actor model primarily increases variance but also adds a small amount of bias towards earlier responses. Due to scalar variability, smaller response magnitudes result in lower production variability. This figure depicts data simulated for the identity context; although we could not measure SMN in the identity context directly, we presume that SMN exists for all behaviors involving a sensorimotor transformation (see **Discussion**).

**Supplementary Figure 4.**
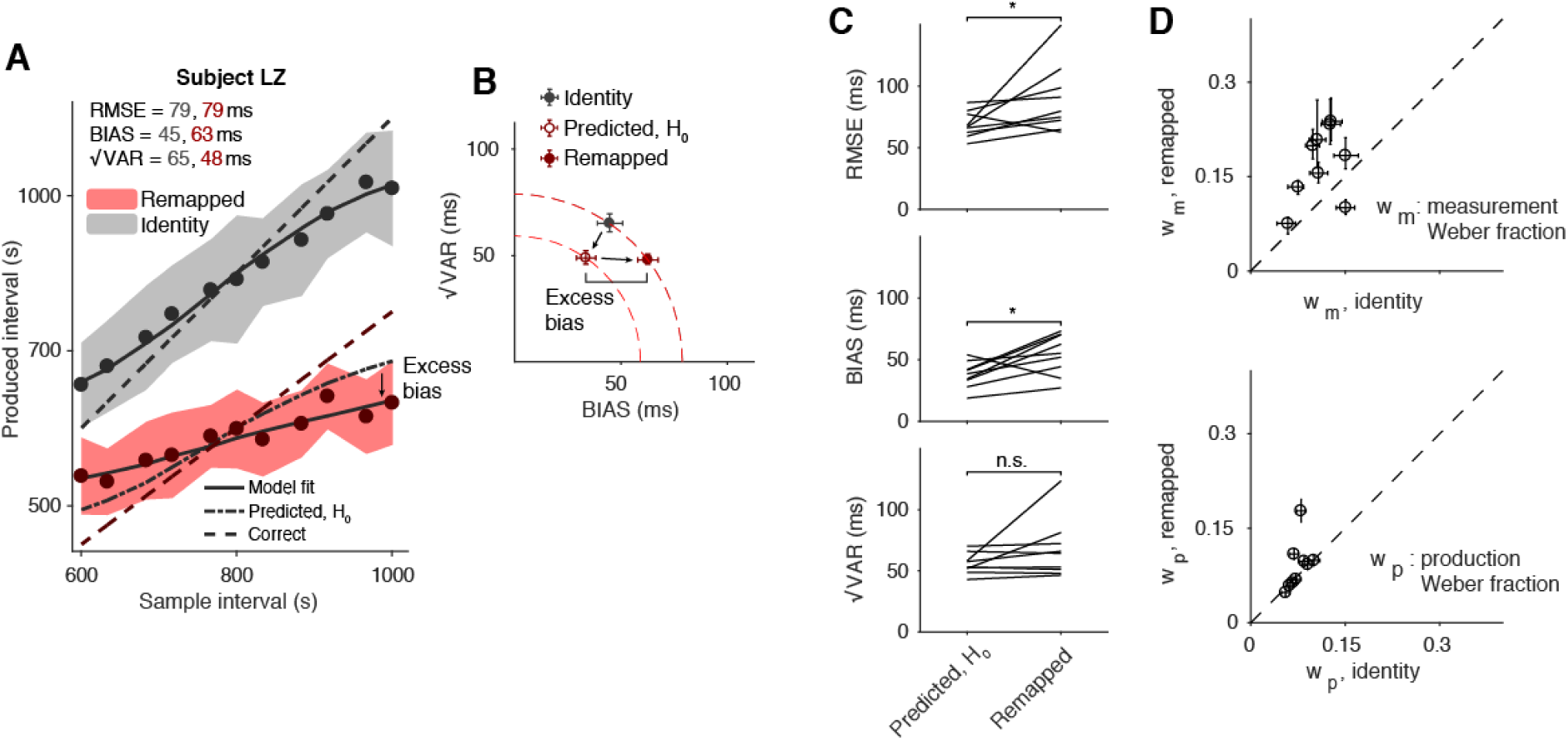
Time interval reproduction, gain = 0.75. An alternative explanation for increased bias in the remapped context (gain = 1.5) which does not involve late inference is that subjects did not follow task instructions, adding a constant duration to their responses rather than multiplying. To investigate this possibility, we designed a control experiment in which the gain factor was 0.75 rather than 1.5. In this case, if subjects added a constant (negative) offset, *BIAS* and fit *w*_*m*_ values should decrease. **A**. Performance of an example subject in the identity (gray) and remapped (red) contexts. Filled circles and shaded regions indicate mean response times ± one standard deviation; dashed lines represent correct intervals. Solid lines represent the mean responses of a Bayesian observer model (see Methods) fit to the subject’s data separately for the two contexts; the dash-dot line in the *g* = 0.75 context represents the model’s behavior using parameters fit from the identity context. This simulation corresponds to the null prediction of no additional SMN in the remapped context. As was the case for *g* = 1.5, excess bias towards the mean in the remapped relative to the identity context is apparent. **B.** *√VAR* vs. *BIAS* for the two contexts (gray for identity, and dark red for remapped), as well as the prediction for the remapped context assuming no additional SMN (empty circle). This prediction underestimates *RMSE* indicating larger SMN for the remapped context. The increased *RMSE* was entirely due to an increase in BIAS (“excess bias”). Dashed quarter circles illustrate combinations of *BIAS* vs. *√VAR* giving rise to equal *RMSE*; error bars are computed from the standard deviation of bootstrapped estimates (n = 1000). **C.** Comparison of *RMSE*, *BIAS*, and *√VAR* for nine subjects. On the left are values predicted in the remapped context (i.e. multiplied by the gain of 0.75), and on the right are the actual values observed from behavior in the remapped context (*: p < 0.05). **D**. Parameters of observer model fits. We fit the an observer model with data of individual subjects independently for the two contexts. Error bars represent 95% confidence intervals estimated using a bootstrap procedure (n = 1000). The results of this experiment suggest that response shifting strategy does not account for increased bias in the remapped contexts in the timing task.

---

**Supplementary Table 1.** Subject performance and observer model parameters for the timing task in the identity and remapped contexts, along with performance predicted for the remapped context under the prediction of no additional SMN. All values are expressed in milliseconds, except for *w*_*m*_ and *w*_*p*_, which are unitless.

| Subject | Identity context |  |  |  |  |  | No additional SMN prediction |  |  | Remapped context |  |  |  |  |  |
| --- | --- | --- | --- | --- | --- | --- | --- | --- | --- | --- | --- | --- | --- | --- | --- |
| | RMSE | BIAS | $\sqrt{\text{VAR}}$ | $w_m$ | $w_p$ | offset | RMSE | BIAS | $\sqrt{\text{VAR}}$ | RMSE | BIAS | $\sqrt{\text{VAR}}$ | $w_m$ | $w_p$ | offset |
| CK | 71 | 37 | 57 | 0.08 | 0.06 | 20 | 106 | 56 | 86 | 136 | 86 | 94 | 0.12 | 0.05 | 32 |
| JM | 103 | 72 | 71 | 0.15 | 0.07 | -22 | 154 | 108 | 106 | 140 | 78 | 108 | 0.11 | 0.07 | -32 |
| MW | 89 | 56 | 69 | 0.13 | 0.07 | -1 | 134 | 84 | 104 | 171 | 128 | 112 | 0.21 | 0.07 | -12 |
| RA | 91 | 46 | 78 | 0.11 | 0.08 | -8 | 136 | 69 | 116 | 212 | 151 | 130 | 0.30 | 0.11 | -54 |
| SMP | 83 | 25 | 77 | 0.06 | 0.08 | -18 | 125 | 38 | 115 | 196 | 121 | 146 | 0.18 | 0.11 | -31 |
| LZ | 79 | 45 | 65 | 0.10 | 0.06 | -6 | 119 | 67 | 98 | 178 | 137 | 112 | 0.24 | 0.08 | -13 |
| DS | 116 | 66 | 94 | 0.15 | 0.10 | -16 | 173 | 98 | 140 | 202 | 136 | 147 | 0.23 | 0.11 | -21 |
| BG | 88 | 52 | 70 | 0.11 | 0.07 | 15 | 132 | 77 | 105 | 191 | 151 | 114 | 0.29 | 0.09 | -25 |

**Supplementary Table 2.** Fit parameters for the five models in the timing task. All values are unitless.

| Subject | Equal-SMN |  |  | Early-Inference |  |  |  | Late-Inference |  |  |  | Late-Ignore-SMN |  |  |  | Observer-Actor |  |  |  |
| --- | --- | --- | --- | --- | --- | --- | --- | --- | --- | --- | --- | --- | --- | --- | --- | --- | --- | --- | --- |
| | $w_m$ | $w_p$ | BIC | $w_m$ | $w_p$ | $w_t$ | BIC | $w_m$ | $w_p$ | $w_t$ | BIC | $w_m$ | $w_p$ | $w_t$ | BIC | $w_m$ | $w_p$ | $w_t$ | BIC |
| CK | 0.10 | 0.06 | -2673 | 0.10 | 0.05 | 0.02 | -2667 | 0.08 | 0.05 | 0.09 | -2705 | 0.10 | 0.06 | 0.00 | -2666 | 0.10 | 0.05 | 0.02 | -2669 |
| JM | 0.13 | 0.07 | -2317 | 0.13 | 0.07 | 0.03 | -2312 | 0.13 | 0.07 | 0.00 | -2310 | 0.13 | 0.07 | 0.00 | -2310 | 0.13 | 0.07 | 0.03 | -2312 |
| MW | 0.16 | 0.07 | -2265 | 0.16 | 0.07 | 0.00 | -2258 | 0.13 | 0.07 | 0.16 | -2308 | 0.16 | 0.07 | 0.00 | -2258 | 0.16 | 0.07 | 0.00 | -2259 |
| RA | 0.17 | 0.09 | -1850 | 0.16 | 0.08 | 0.06 | -1852 | 0.10 | 0.09 | 0.25 | -1958 | 0.16 | 0.09 | 0.10 | -1847 | 0.16 | 0.08 | 0.06 | -1855 |
| SMP | 0.12 | 0.10 | -1778 | 0.10 | 0.08 | 0.08 | -1802 | 0.05 | 0.10 | 0.17 | -1915 | 0.09 | 0.09 | 0.13 | -1812 | 0.10 | 0.08 | 0.09 | -1807 |
| LZ | 0.15 | 0.07 | -2237 | 0.15 | 0.07 | 0.05 | -2237 | 0.10 | 0.07 | 0.20 | -2362 | 0.15 | 0.07 | 0.02 | -2230 | 0.15 | 0.07 | 0.05 | -2240 |
| DS | 0.18 | 0.11 | -1655 | 0.18 | 0.10 | 0.04 | -1649 | 0.15 | 0.11 | 0.16 | -1663 | 0.18 | 0.10 | 0.09 | -1649 | 0.18 | 0.10 | 0.04 | -1650 |
| BG | 0.17 | 0.08 | -2094 | 0.16 | 0.08 | 0.04 | -2089 | 0.11 | 0.08 | 0.24 | -2206 | 0.17 | 0.08 | 0.00 | -2087 | 0.16 | 0.08 | 0.04 | -2092 |
| Total |  |  | -16803 |  |  |  | -16783 |  |  |  | -17344 |  |  |  | -16776 |  |  |  | -16800 |

**Supplementary Table 3.** Subject performance and observer model parameters for the length task in the identity and remapped contexts, along with performance predicted for the remapped context under the prediction of no additional SMN. All values are expressed in degrees visual angle, except for *w*_*m*_ and *w*_*p*_, which are unitless.

|  | Identity context |  |  |  |  |  |  | No additional SMN prediction |  |  | Remapped context |  |  |  |  |  |  |
| --- | --- | --- | --- | --- | --- | --- | --- | --- | --- | --- | --- | --- | --- | --- | --- | --- | --- |
| Subject | RMSE | BIAS | $\sqrt{\text{VAR}}$ | $w_m$ | $w_p$ | $\sigma_p$ | offset | RMSE | BIAS | $\sqrt{\text{VAR}}$ | RMSE | BIAS | $\sqrt{\text{VAR}}$ | $w_m$ | $w_p$ | $\sigma_p$ | offset |
| EG | 0.94 | 0.13 | 0.93 | 0.00 | 0.07 | 0.00 | 0.17 | 1.38 | 0.15 | 1.38 | 1.60 | 0.52 | 1.51 | 0.06 | 0.06 | 0.66 | -0.24 |
| IG | 1.77 | 0.51 | 1.69 | 0.08 | 0.06 | 1.30 | 1.15 | 2.13 | 0.77 | 1.99 | 2.63 | 1.23 | 2.32 | 0.12 | 0.11 | 0.00 | -0.44 |
| JK | 1.38 | 0.35 | 1.33 | 0.06 | 0.06 | 0.89 | 0.90 | 1.77 | 0.52 | 1.69 | 1.84 | 0.86 | 1.62 | 0.09 | 0.01 | 1.28 | -0.29 |
| RC | 1.28 | 0.24 | 1.25 | 0.04 | 0.06 | 0.90 | 0.69 | 1.58 | 0.35 | 1.54 | 1.79 | 0.83 | 1.58 | 0.08 | 0.07 | 0.00 | -0.27 |
| DS | 1.43 | 0.49 | 1.35 | 0.07 | 0.06 | 0.91 | 0.56 | 1.88 | 0.72 | 1.73 | 2.34 | 0.76 | 2.21 | 0.08 | 0.10 | 0.56 | -0.33 |
| SMP | 1.75 | 0.57 | 1.65 | 0.07 | 0.01 | 1.55 | 0.59 | 1.97 | 0.75 | 1.82 | 2.33 | 0.99 | 2.11 | 0.09 | 0.00 | 1.85 | -0.22 |
| TT | 0.62 | 0.07 | 0.62 | 0.00 | 0.05 | 0.17 | -0.10 | 0.92 | 0.10 | 0.91 | 1.62 | 0.31 | 1.59 | 0.03 | 0.08 | 0.00 | -0.22 |

**Supplementary Table 4.** Fit parameters the for five models in the length task. All values are unitless except *σ*_*p*_, which is in units of visual degrees.

|  | Equal-SMN |  |  |  | Early-Inference |  |  |  |  | Late-Inference |  |  |  |  | Late-Ignore-SMN |  |  |  |  | Observer-Actor |  |  |  |  |
| --- | --- | --- | --- | --- | --- | --- | --- | --- | --- | --- | --- | --- | --- | --- | --- | --- | --- | --- | --- | --- | --- | --- | --- | --- |
| Subject | $w_m$ | $w_p$ | $\sigma_p$ | BIC | $w_m$ | $w_p$ | $\sigma_p$ | $w_t$ | BIC | $w_m$ | $w_p$ | $\sigma_p$ | $w_t$ | BIC | $w_m$ | $w_p$ | $\sigma_p$ | $w_t$ | BIC | $w_m$ | $w_p$ | $\sigma_p$ | $w_t$ | BIC |
| EG | 0.03 | 0.07 | 0.00 | 5720 | 0.02 | 0.07 | 0.00 | 0.16 | 5905 | 0.00 | 0.07 | 0.32 | 0.06 | 5608 | 0.01 | 0.07 | 0.34 | 0.06 | 5695 | 0.02 | 0.06 | 0.39 | 0.21 | 5883 |
| IG | 0.10 | 0.10 | 0.66 | 4988 | 0.09 | 0.10 | 0.83 | 0.15 | 5154 | 0.08 | 0.10 | 0.77 | 0.09 | 4983 | 0.09 | 0.09 | 1.00 | 0.09 | 4993 | 0.09 | 0.08 | 1.03 | 0.23 | 5152 |
| JK | 0.07 | 0.04 | 1.08 | 6485 | 0.07 | 0.04 | 1.08 | 0.08 | 6728 | 0.06 | 0.03 | 1.15 | 0.06 | 6474 | 0.07 | 0.04 | 1.08 | 0.00 | 6492 | 0.06 | 0.02 | 1.18 | 0.17 | 6724 |
| RC | 0.06 | 0.05 | 0.88 | 6287 | 0.06 | 0.06 | 0.85 | 0.00 | 6529 | 0.04 | 0.04 | 1.01 | 0.07 | 6253 | 0.06 | 0.05 | 0.88 | 0.00 | 6294 | 0.05 | 0.04 | 0.96 | 0.14 | 6527 |
| DS | 0.08 | 0.10 | 0.00 | 6967 | 0.07 | 0.07 | 0.78 | 0.30 | 7199 | 0.07 | 0.10 | 0.00 | 0.03 | 6974 | 0.07 | 0.10 | 0.17 | 0.07 | 6968 | 0.06 | 0.08 | 0.73 | 0.29 | 7201 |
| SMP | 0.09 | 0.07 | 1.33 | 4880 | 0.07 | 0.00 | 1.56 | 0.26 | 5036 | 0.07 | 0.06 | 1.37 | 0.06 | 4883 | 0.08 | 0.06 | 1.40 | 0.04 | 4887 | 0.07 | 0.00 | 1.56 | 0.28 | 5033 |
| TT | 0.00 | 0.07 | 0.00 | 5163 | 0.02 | 0.05 | 0.00 | 0.29 | 5224 | 0.00 | 0.06 | 0.00 | 0.05 | 5077 | 0.01 | 0.05 | 0.01 | 0.07 | 5054 | 0.01 | 0.05 | 0.01 | 0.29 | 5204 |
| Total |  |  |  | 40559 |  |  |  |  | 41857 |  |  |  |  | 40334 |  |  |  |  | 40466 |  |  |  |  | 41805 |

